# Genome-wide identification and functional analysis of circRNAs in *Zea mays*

**DOI:** 10.1101/384693

**Authors:** Baihua Tang, Zhiqiang Hao, Yanfeng Zhu, Hua Zhang, Guanglin Li

## Abstract

Circular RNAs (circRNAs) are a class of endogenous noncoding RNAs, which increasingly drawn researchers’ attention in recent years as their importance in regulating gene expression at the transcriptional and post-transcriptional levels. With the development of high-throughput sequencing and bioinformatics, circRNAs have been widely analysed in animals, but the understanding of characteristics and function of circRNAs is limited in plants, especially in maize. Here, 3715 unique circRNAs were predicted in *Zea mays* systematically, and 8 of 12 circRNAs were validated by experiments. By analysing circRNA sequence, the events of alternative circularization phenomenon were found prevailed in maize. By comparing circRNAs in different species, it showed that part circRNAs are conserved across species, for example, there are 273 circRNAs conserved between maize and rice. Although most of the circRNAs have low expression levels, we found 213 differential expressed circRNAs responding to heat, cold, or drought, and 1782 tissue-specific expressed circRNAs. The results showed that those circRNAs may have potential biological functions in specific situations. Finally, two different methods were used to search circRNA functions, which were based on circRNAs originated from protein-coding genes and circRNAs as miRNA decoys. 346 circRNAs could act as miRNA decoys, which might modulate the effects of multiple molecular functions, including binding, catalytic activity, oxidoreductase activity, and transmembrane transporter activity. Maize circRNAs were identified, classified and characterized systematically. We also explored circRNA functions, suggesting that circRNAs are involved in multiple molecular processes and play important roles in regulating of gene expression. Our results provide a rich resource for further study of maize circRNAs.

## Introduction

Circular RNAs (circRNAs) are a class of endogenous noncoding RNA molecules formed by backsplicing [1-5]. Although circRNAs or circular isoforms (e.g., muscleblind gene, sodium transporter NCX1, the rat cytochrome P450 2C24 gene, ETS and the cytochrome P450 2C18 genes) have been discovered in *Drosophila*, mice, and humans many years [6-10], they have increasingly drawn researchers’ attention in recent years for their important roles in regulating gene expression.

Unlike mRNA, circRNA transcripts usually lack of the 5’ cap and 3’ poly(A) tail, and the majority of circRNAs are expressed at low levels compared with linear RNAs [1, 3, 5, 11]. However, some specific circRNAs have prominent expression compared to their corresponding linear isoforms, for instance, two antisense transcripts of CDR1as and cANRIL [2, 12]. Different circRNAs are often expressed in specific tissues, cell types or developmental stages [2, 5, 13, 14], and particular time courses, suggesting that circRNAs exhibit spatiotemporal-specific expression patterns [15]. Additionally, certain circRNAs showed evolutionary conservation between humans and mice [2, 3], play roles in regulating gene expression, and are often present in some diseases [12, 16-19]. Intriguingly, circRNAs can act as miRNA or RNA binding protein (RBP) sponges, which sequester miRNAs away from their mRNA targets [2, 14, 20-22], for example, CDR1as and Sry as miRNA sponges [2, 23], inferring circRNA regulatory functions in the genetic network [11, 13]. With development of high-throughput sequencing, the methods identifying circRNAs have been developed [1-5, 13], and a number of circRNAs have been identified in animals. However, plant circRNAs are still underappreciated with the exception of those in thale cress (*Arabidopsis thaliana*), rice (*Oryza sativa*), tomato (*Solanum lycopersicum*), barley (*Hordeum vulgare*), maize (Zea *mays L*.) and trifoliate orange (*Poncirus trifoliata L. Raf*.) [11, 15, 24-30].

Maize (*Zea mays L*.) is one of the most important crops worldwide and serves as model organism in biological research. With the development of high-throughput sequencing technology, more and more data have been produced, now it is possible for us to study circRNAs in maize systematically. In this article, maize circRNAs were firstly identified from multiple resources. Then circRNA characteristics, such as genomic distribution, alternative circularization, conservation and expression patterns, were analyzed. Whether CircRNAs act as miRNA decoys to mediate the regulation of gene expression in maize or not were analysed. Finally, maize circRNA functions were inferred in our study. The discovery of maize circRNAs enriches the repositories of plant circRNAs.

## Results

### Identification and Classification of maize circRNAs

In order to identify circRNAs in *Zea mays*, transcriptome data were firstly collected from 5 resources (Table 1), then find_circ, one of methods widely used in circRNA prediction [2], was carried out for the genome-wide identification of circRNAs (S1 Fig), finally we predicted 7011 circRNAs totally. After merging the circRNAs with same loci, 3715 unique circRNAs candidates were used for further analysis (S1 Table).

**Table 1.**
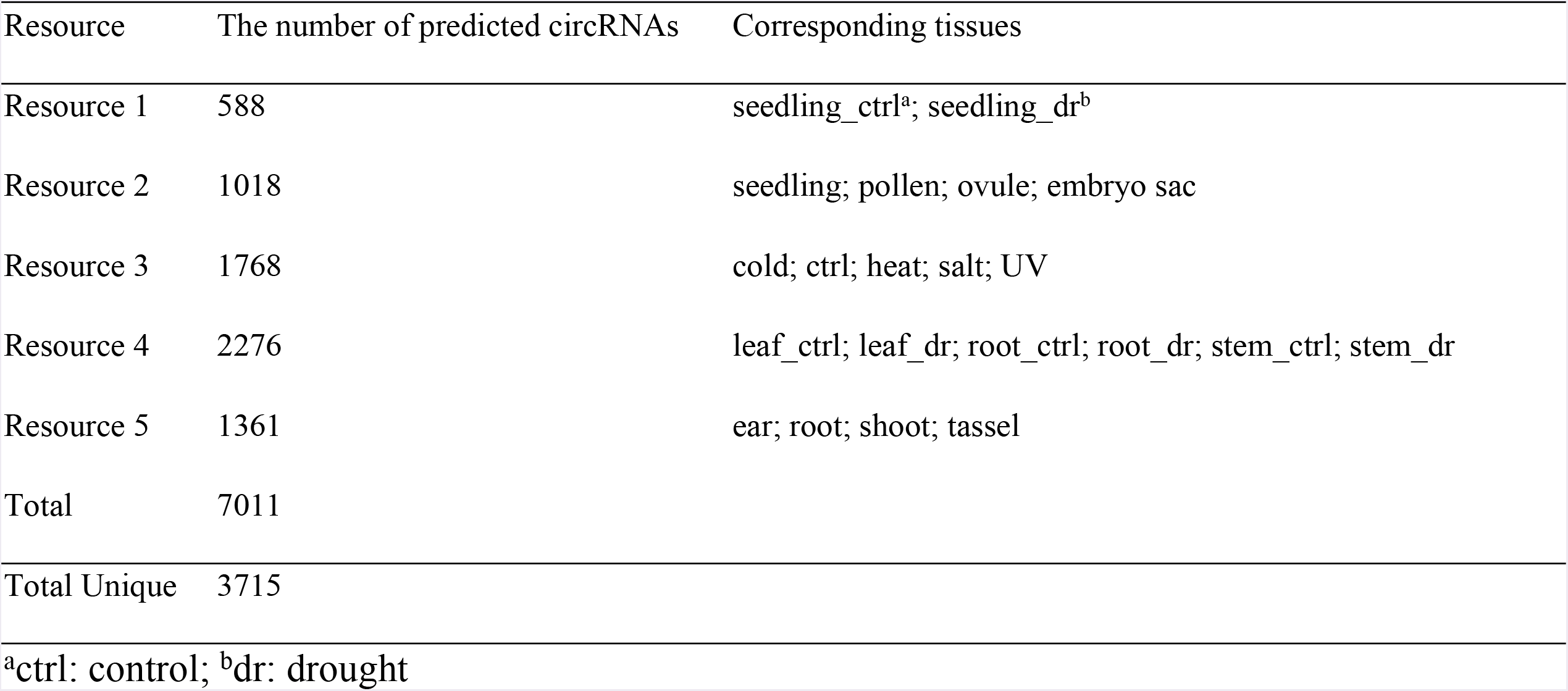
The number of predicted circRNAs from five resources.

All predicted circRNAs can be classified into three different groups based on positional relationship between the circRNA and their related gene: “circRNAs within genes”, “circRNAs overlap with genes” and “intergenic circRNAs”, with proportions of 47.6%, 19.7%, and 32.7%, respectively (S1 Table). In addition, based on the positional relationship between the circRNA and their related exons, circRNAs can be divided into exonic circRNAs (ecircRNAs) and non-ecircRNAs. In maize circRNAs, there are 2007 ecircRNAs, the backsplice sites of which are located in the CDS-CDS, CDS-3’ UTR, CDS-5’ UTR, 3’ UTR-3’ UTR, 3’ UTR-5’ UTR, and 5’ UTR-5’ UTR. EcircRNAs account for 54% of the total circRNAs, which constitute the main part of the circRNAs (Fig 1A).

**Fig 1.**
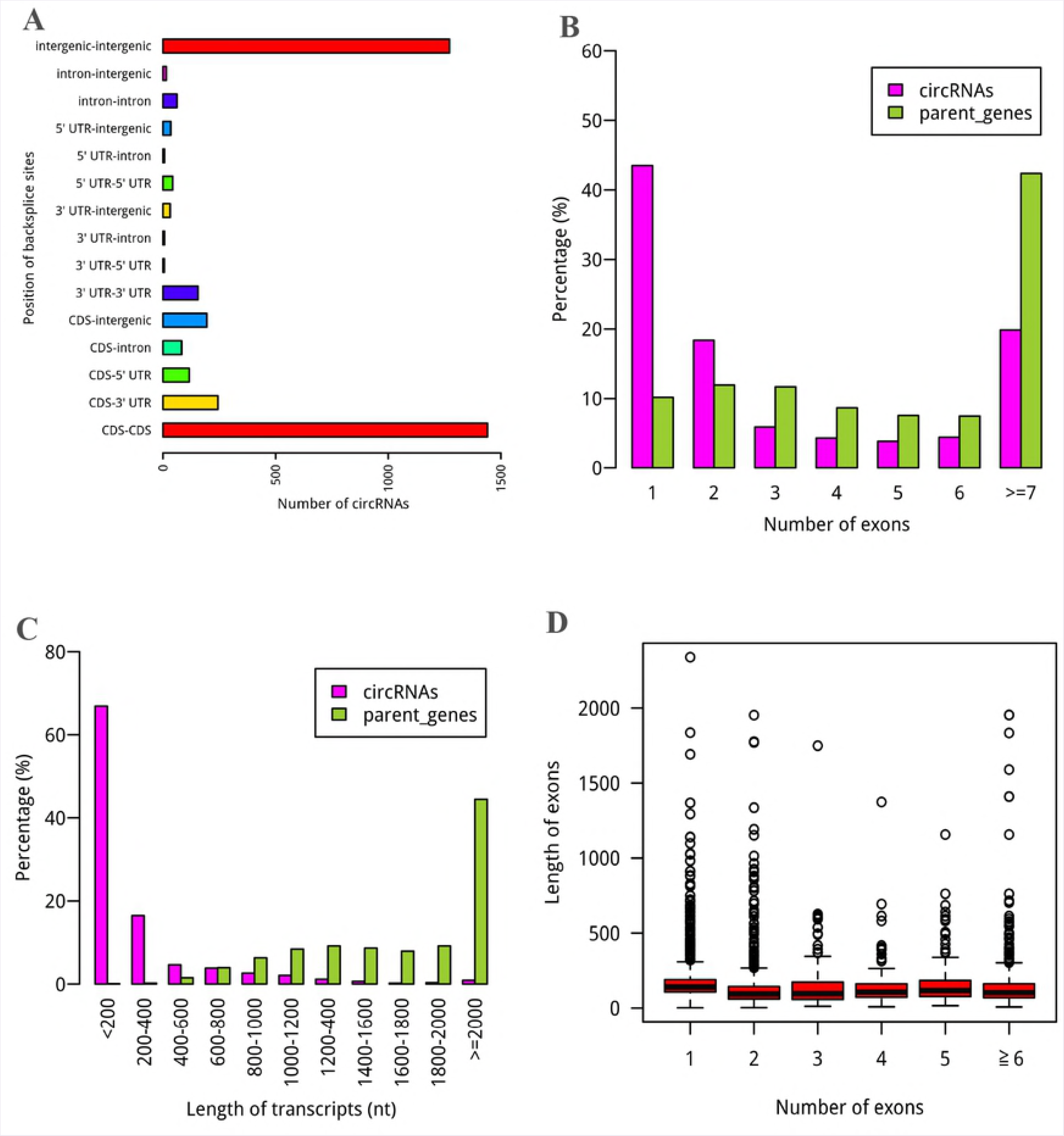
General feature of the circRNA candidates. (A) Genomic locations of the circRNAs. The backsplice site was predominantly located at CDS-only and intergenic-only. (B) Exon number distributions of circRNAs and their parent genes. (C) Length distributions of circRNAs and their parent genes. (D) Length distributions of back-spliced exons corresponding to different numbers of exons.

### Characteristics of circRNAs in maize

When comparing numbers of circRNAs with backsplice sites located in different regions (Fig 1A), we found more circRNAs at 3’ UTR-3’ UTR than at 5’ UTR-5’ UTR, and the same case happened between CDS-3’ UTR and CDS-5’ UTR. The results showed that circRNAs are more frequency in 3’ UTR than in 5’ UTR. Interestingly, some circRNAs also found in introns. The un-random distributions of circRNAs in genome suggest that they have biological functions. Majority of circRNAs contained 1 to 4 exons less than their corresponding parent genes, most of which had more than 6 exons (Fig 1B). Likewise, the comparison of transcript lengths between the circRNAs and their parent genes displayed circRNAs with shorter lengths (*P* < 2.2e-16, Wilcoxon rank-sum test) (Fig 1C). The two results are in accordance with previous reports [15, 25]. Interestingly, circularized exons from single exon circRNAs have a slightly longer length than exons from circRNAs with multiple circularized exons (Fig 1D), suggesting that longer exons may facilitate circularization [3, 31, 32].

The intron length distributions were also compared between the exonic circRNAs and linear genes. We found that introns flanking circularized exon were significantly longer than that of the linear genes (*P* < 2.2e-16, Wilcoxon rank sum test) (Fig 2A), as was demonstrated in previous studies [1, 3, 15, 33, 34]. This observation suggests that longer flanking introns may promote exon circularization. However, determining whether the existence of these structures is necessary for circRNA formation requires further investigation.

**Fig 2.**
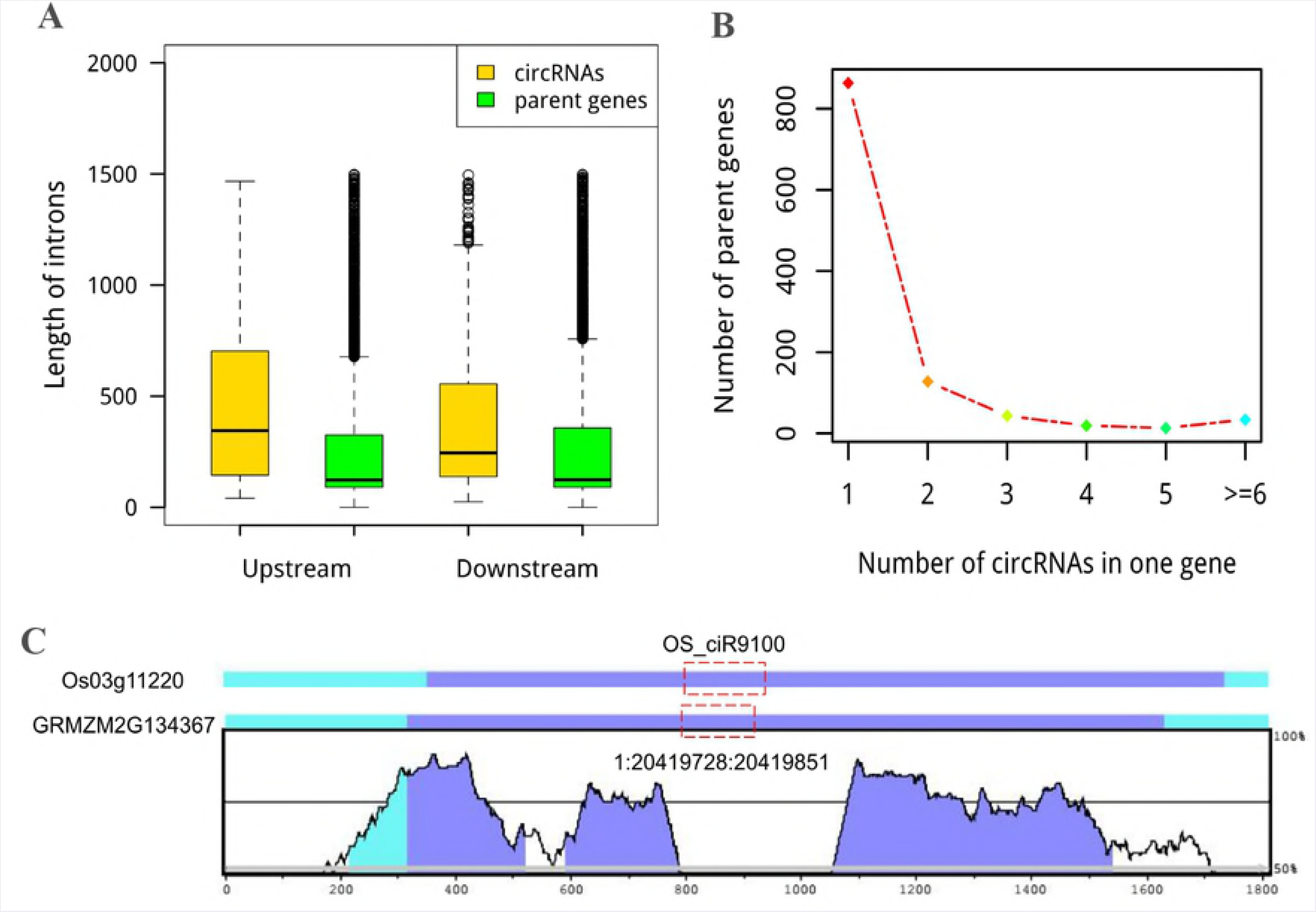
The genomic character and conservation of circRNAs. (A) Mean length distribution of upstream and downstream intron flanking circularized exons in *Z.mays*. Longer intron bracketing circularized exons was observed than the parent gene. (B) The number of genes (y-axis) that generated different numbers of circRNAs (x-axis). (C) An example of sequence conservation analysis of circRNAs between *O. sativa* and *Z. mays*. Y-axis, levels of conservation based on genomic sequence similarity. The locations of circRNAs in their respective parent genes are marked with a red box. Conserved regions are labelled in different colours by VISTA according to the annotations (exons in blue shadow, UTR in cyan shadow and CNS (non-coding sequences) in red shadow).

Because different forms of linear mRNAs are produced during the alternative splicing (AS) process in *Z. mays* [35], we hypothesized that alternative circularization (multiple circRNAs produced from one gene) happened on circRNAs. Based on our results, 905 exon circularization events from 237 parent genes were discovered, with two circRNA isoforms from one gene making up larger proportion (54.0%), three different circRNAs accounting for 18.1%, four for 8.0%, five for 5.5%, and the rest of the genes possessed at least six circRNAs (Fig 2B and S2 Table). Among these alternative circularization events, one parent gene possessed the most 32 circRNAs based on annotated information. Examples of alternative circularization are shown in S2A Fig. Obviously, circRNAs comprise single or multiple exons with or without known exon boundaries from the same gene. Most circRNAs (1431/1768) used new exon boundary, only 90 circRNAs had two annotated exon boundaries, and 247 circRNAs with one known boundary (S2 Table). As a result, ubiquitous alternative circularization increased the complexity of circRNAs in *Z. mays*.

### Conservation of circRNAs between maize and other species

To evaluate circRNA conservation among different species, we firstly compared circRNAs in *Z. mays* with circRNAs in *O. Sativa* and *A. thaliana* [15]. 291 orthologous gene pairs were found in parent genes of circRNA, and a total of 232 circRNAs were found conserved between *Z. mays* and *O. sativa* (Fig 2C and S3A Table), which comprised 14.5% circRNAs parent genes in *Z. mays* higher than the proportion in *O. sativa* (12.2%) in a previous publication [15]. For parent genes in *Z. mays* and *A. thaliana*, 109 orthologous gene pairs were found, and a total of 122 circRNAs were conserved between *Z. mays* and *A. Thaliana*. In addition, the ratio of orthologous genes to the total parent genes (10.9%) in *Z. mays* was slightly lower than that in *A. thaliana* (14.5%) [15] (S2B Fig and S3B Table). When merging the conservation information of maize circRNAs in the above two parts, totally 307 conserved maize circRNAs can be identified, in which 47 circRNAs are conserved among maize, rice and arabidopsis. In addition, when comparing maize circRNAs with published circRNAs from tomato and soybean, 115 and 149 maize circRNAs are conserved, respectively (S3C and S3D Table).

### Validation of circular RNAs in maize

To confirm our identification of maize circRNAs, twelve randomly selected circRNAs from the highly expressed maize circRNAs were used in experimental validation. Divergent primers (S4 Table) were firstly designed for each circRNA, then reverse transcription PCR were used to amplify both cDNA and genomic DNA, respectively. Theoretically, it was expected that positive and negative results of amplification would be obtained for cDNA and genomic DNA, respectively. As a control, convergent primers were also designed for each circRNA and used to amplify the linear mRNAs in verification. The amplified PCR products using divergent primers were sequenced to confirm the presence of the back spliced junctions of circRNAs. As a result, 8 of the 12 circRNAs were validated (Fig 3).

**Fig 3.**
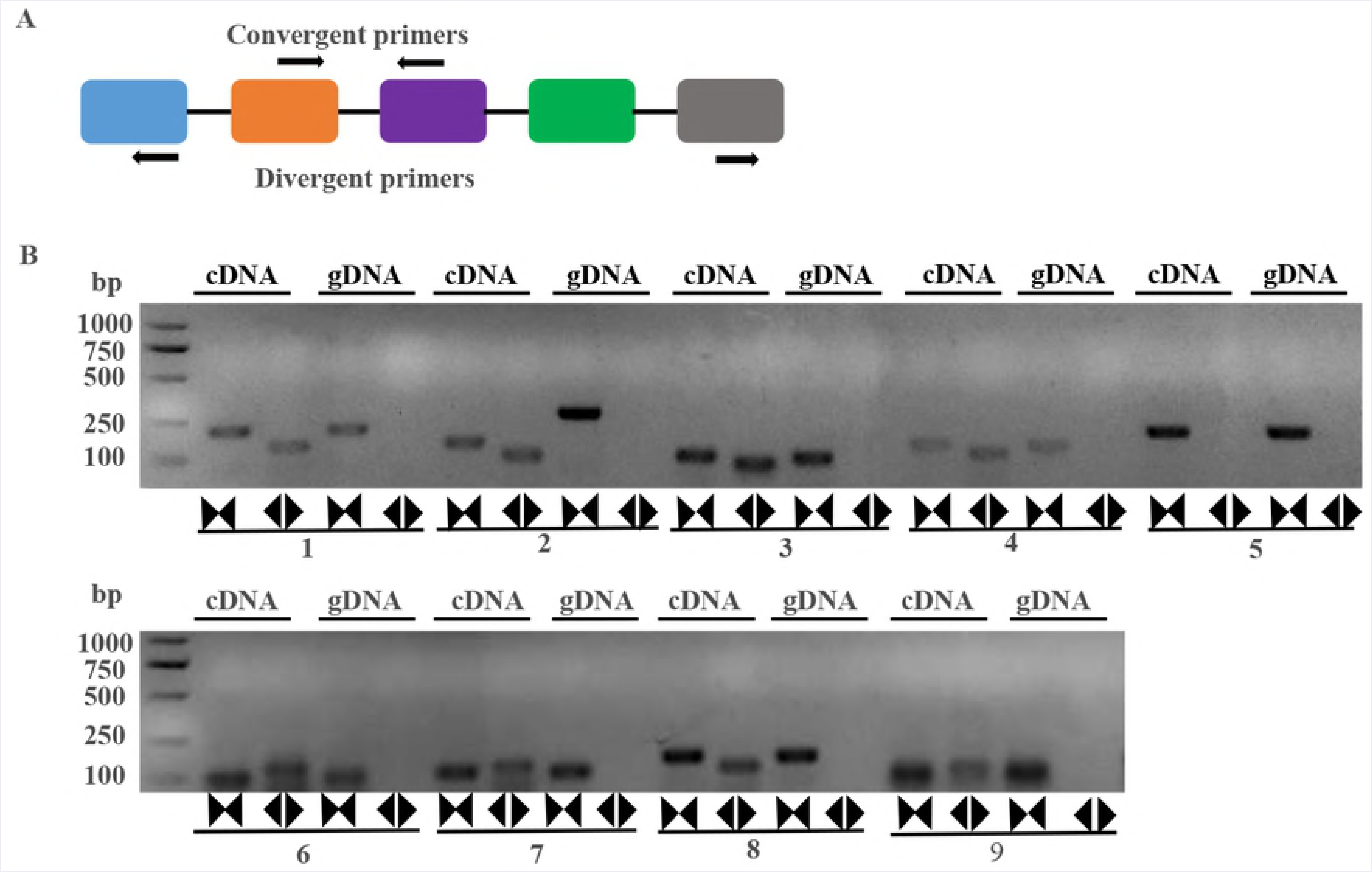
Validation of maize circRNAs. Validation of circular RNAs (circRNAs) by PCR amplifications with divergent primers in maize. (A) Divergent and convergent primers for amplification of circRNA and linear RNA are shown in a model, respectively. (B) A set of divergent primers (black back-to-back triangle pairs) successfully amplified eight circRNAs in cDNA. A set of convergent primers (black opposing triangle pairs) could work on both cDNA and genomic DNA. Note: 1: seedling_test_circ_000962, 2: seedling_test_circ_001425, 3: seedling_test_ circ_000119, 4: seedlingd_test_circ_002114, 5: Actin, 6: seedlingc_test_circ_000623, 7: seedlingd_ test_circ_000813, 8: seedling_test_circ_001637, 9: control_test_circ_004099.

### Expression pattern of maize circRNAs

Twenty-one samples were used to measure the expression abundance of the circular RNAs. The result of two different measurements both showed that expression levels of maize circRNAs were primarily concentrated in 0-0.5 ratio or RPM bin (Fig 4A and S3 Fig). Apparently, the quantitative distribution of circRNAs in the various intervals was more dispersed than the distribution of their parent genes (Fig 4A and S3 Fig). Interestingly, the number of circRNAs with outstanding expression level in the sample of heat was more than other conditions (Fig 4B). Additionally, the Pearson correlation coefficient values were calculated to examine whether the circRNAs were capable of regulating the expression of their parent genes as reported in animals [36, 37]. We found 791 significantly positively correlated pairs and 2 significantly negatively correlated pairs amongst 1818 total pairs of circRNAs and their parent genes (*P* < 0.05), suggesting their correlation for gene expression regulation (S5 Table).

**Fig 4.**
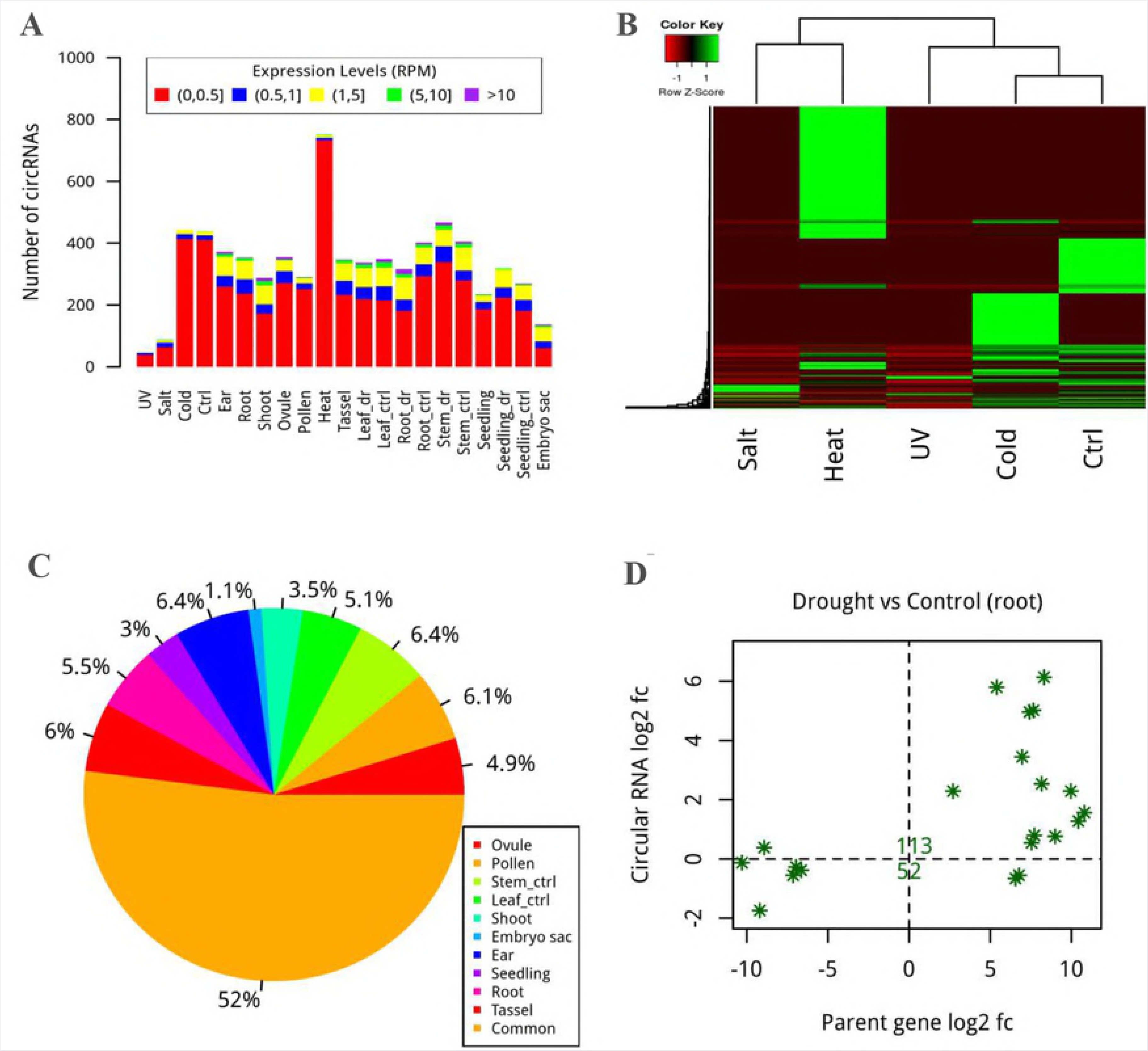
Expression pattern of circRNAs. (A) Expression level of circRNAs in different scopes. CircRNAs were usually expressed at lower levels, which may explain their difficult detection because they lack a poly(A) tail. ctrl and dr indicate control and drought, respectively. (B) The degree of circRNA expression under five different treatments was shown by heatmap. More circRNAs were present under heat stress. (C) Pie chart with rainbow colours showing the ratio of circRNAs possessing tissue-specific expression in 10 different tissues. (D) Tendency for differential expression of circRNAs and the overlapping genes was plotted using fold changes between the drought and control in root as the standard. The number of both differentially expression circRNAs and genes is marked.

To investigate whether circRNAs have tissue/developmental stage-specific expression patterns [2, 13], transcriptome data from 10 tissues containing 21 samples were used to calculate the tissue-specific expression levels. The results showed that half of the circRNAs possessed tissue-specific expression characteristics with a tissue-specific index greater than 0.9. The number of these circRNAs (1782) with tissue-specific expression was similar in all tissues with the exception of the embryo sac (Fig 4C and S3C Fig). Most circRNAs (1537) were expressed in single tissue, although a few circRNAs (13) were commonly expressed in all 10 tissues. The tissue-specific expression patterns of the circRNAs indicated that they might have specific spatial functions.

To examine the differential expression levels of the circRNAs, we compared the expression profiles of circRNAs and genes under control and stress conditions in the different transcriptome data groups. We found that 213 circRNAs were differentially expressed, of which 132 and 117 were up-regulated and down-regulated under heat or cold in seedling, and under drought in stem, leaf, or root, respectively (Table 2). CircRNAs were inclined to be up-regulated or down-regulated with their parent genes simultaneously, except few circRNAs with reverse patterns comparing with their parent genes (Fig 4D and S3 Fig).

**Table 2.**
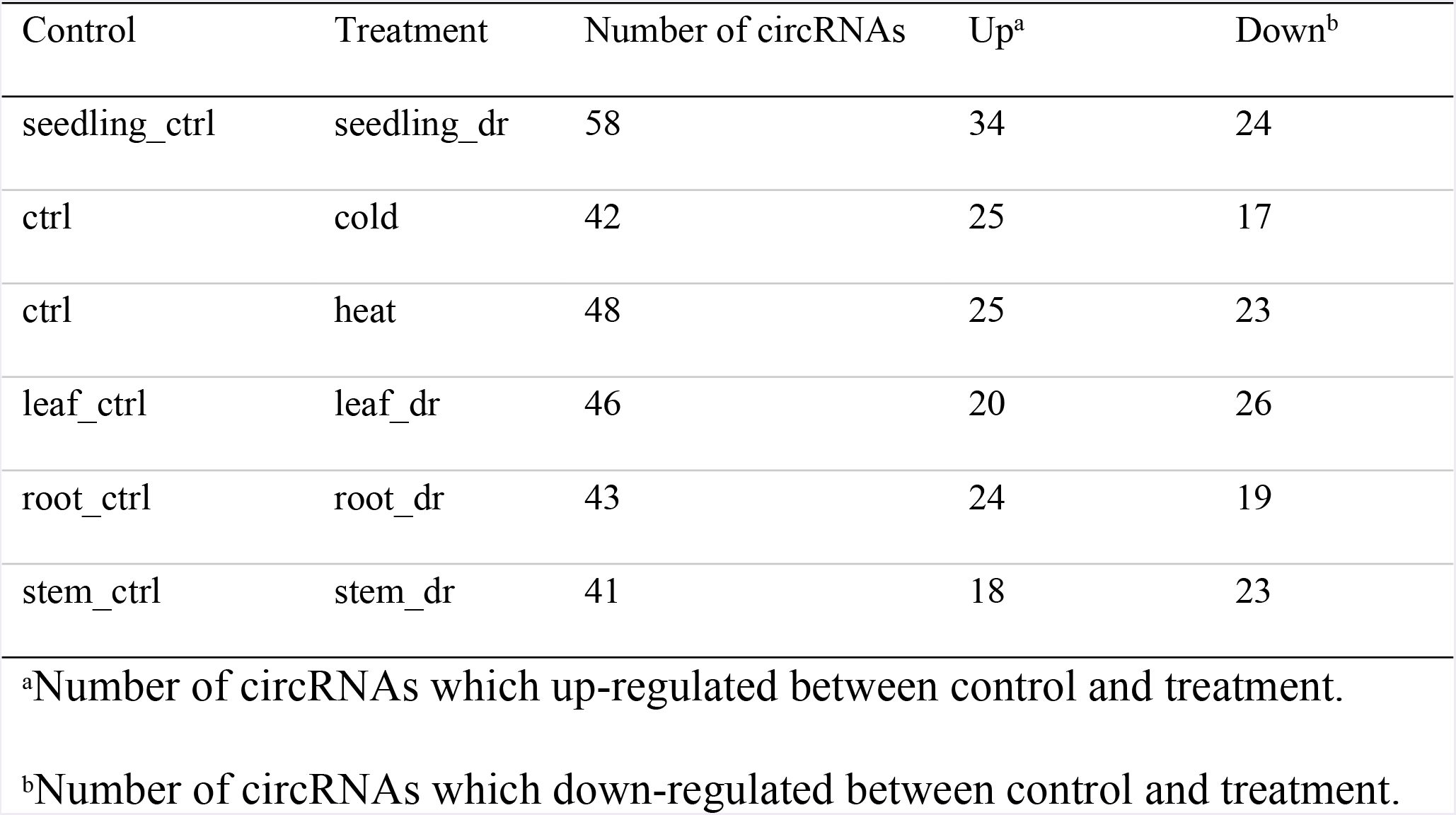
Differential expression of circRNAs between control and treatment.

| Control | Treatment | Number of circRNAs | Up<sup>a</sup> | Down<sup>b</sup> |
| --- | --- | --- | --- | --- |
| seedling_ctrl | seedling_dr | 58 | 34 | 24 |
| ctrl | cold | 42 | 25 | 17 |
| ctrl | heat | 48 | 25 | 23 |
| leaf_ctrl | leaf_dr | 46 | 20 | 26 |
| root_ctrl | root_dr | 43 | 24 | 19 |
| stem_ctrl | stem_dr | 41 | 18 | 23 |

<sup>a</sup>Number of circRNAs which up-regulated between control and treatment.
<sup>b</sup>Number of circRNAs which down-regulated between control and treatment.

### Possible functions of circRNAs

#### Function of circRNAs originated from protein-coding genes

As circRNAs have similar or reverse expression pattern with their parent protein-coding genes, we first performed an enrichment analysis for their parent genes to indirectly infer circRNAs’ roles in maize. We found that 1821 of 3715 circRNAs enriched in 1127 parent genes. The GO enrichment analysis showed that parent genes of these circRNAs were primarily associated with cytoplasm or cytoplasmic parts and two other cellular components (plastid and thylakoid). They are also involved in most of molecular functions including catalytic activity, different enzyme activities (lyase, transferase and methyltransferase) and binding to unfolded proteins and ions. In addition, the parent genes were associated with diverse biological processes, such as response to stress, photosynthesis, and protein folding (S4 Fig). The KEGG results further showed that circRNAs’ parent genes were primarily related to the metabolism of some biomacromolecules (purine metabolism, porphyrin and chlorophyll metabolism, starch and sucrose metabolism, *etc*), amino acids and cell respiration (glycolysis and citrate cycle) (S6 Table).

#### circRNAs acting as miRNA decoys

In order to investigate the function of circRNAs as miRNA decoys in maize, we firstly identified 346 circRNAs which can act as miRNA decoys based on previous methods [38, 39]. Then genome-wide miRNA-circRNA-mRNA networks were constructed to investigate the functions of circRNAs acting as miRNA decoys. The networks were composed of 5269 nodes and 6861 edges; the nodes included 144 miRNAs, 346 circRNAs and 4779 mRNAs (S4 Fig). There were 1314 interactions between these 144 miRNAs and 346 circRNAs acting as miRNA decoys (Fig 5A and S7 Table). We found that the majority of subnets are interconnected in the global regulatory networks (Fig 5B). Interestingly, there were no separate sub-networks between the circRNAs and miRNAs in the miRNA-circRNA-mRNA networks as described above (S4 Fig).

**Fig 5.**
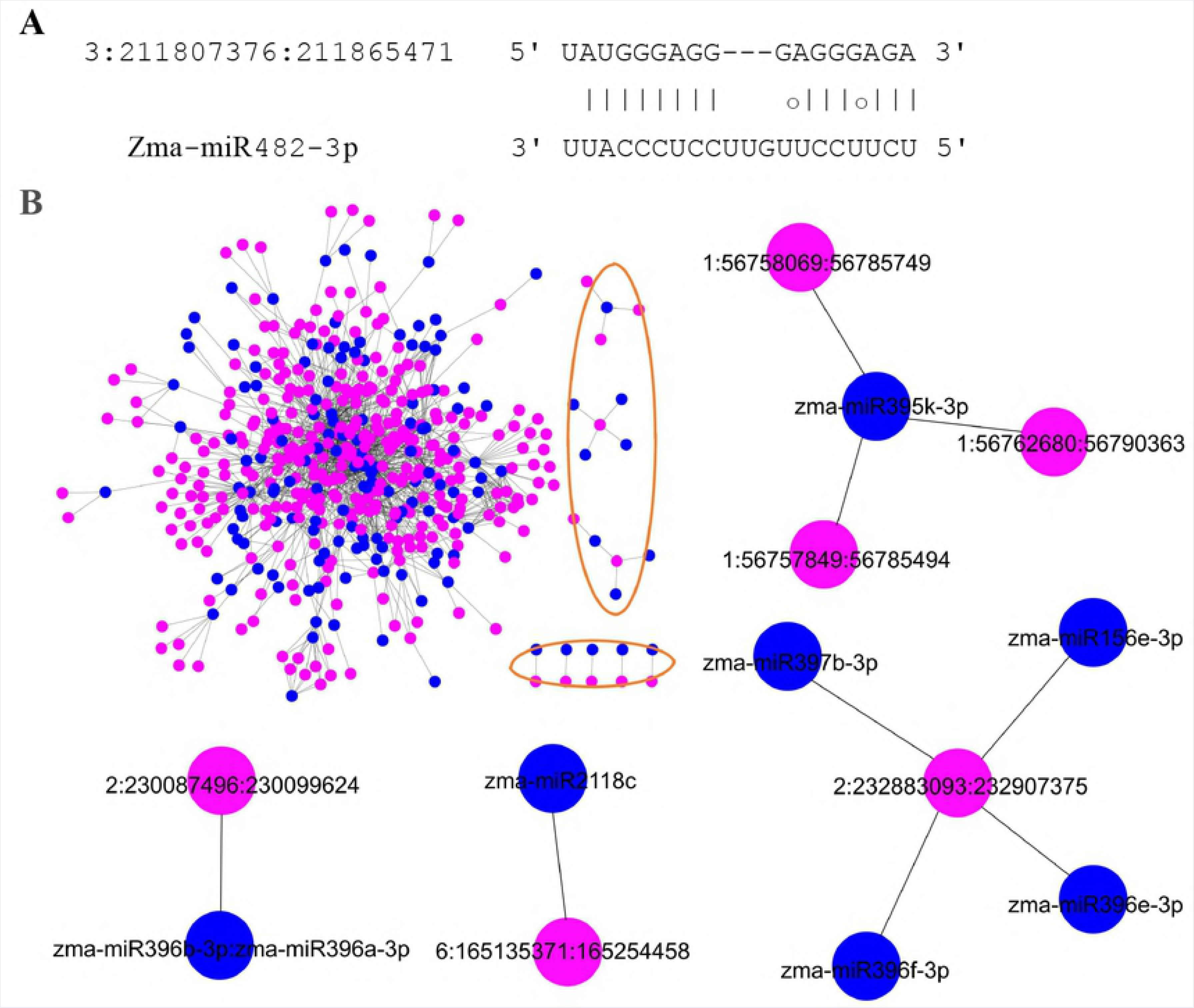
circRNAs acting as miRNA decoys. (A) Binding sites between a circRNA and the corresponding miRNA. (B) The circRNA-miRNA regulatory network. The single network was marked by orange circle. Representative single network were extracted from the integral network. Pink nodes represent circRNAs and blue nodes represent miRNAs. The edges represent connected nodes that exist as a correlation.

Based on the ceRNA hypothesis [40], the function of circRNAs acting as miRNA decoys can be inferred from mRNAs which are in the same subnets with circRNAs. Totally we inferred the functions of 346 circRNAs via 4779 mRNAs (Fig 6). The GO enrichment and functional analysis of the mRNAs suggested that these 346 circRNAs might participate in diverse biological processes, such as multiple metabolic processes, cellular processes, and single organism processes. These circRNAs could also be involved in the formation of cells or cell parts, cytoplasm or cytoplasmic parts, intracellular or intracellular parts and some organelles. Moreover, these circRNAs might modulate the effects of multiple molecular functions, including binding, catalytic activity, oxidoreductase activity, and transmembrane transporter activity. Consequently, circRNAs may play significant roles in various cell locations, metabolic processes, and stress responses (S8 Table). Several pathways about energy metabolism were found, like carbon fixation pathways (25 circRNAs with 7 parent genes, p-value < 0.05), nitrogen metabolism (11 circRNAs with 3 parent genes, p-value < 0.05), starch and surcrose metabolism (22 circRNAs with 17 parent genes, p-value = 0.055) and photosynthesis (1 circRNA with 1 parent gene, p-value = 0.018), showing that circRNAs play important roles in plant yield (S6B Table).

**Fig 6.**
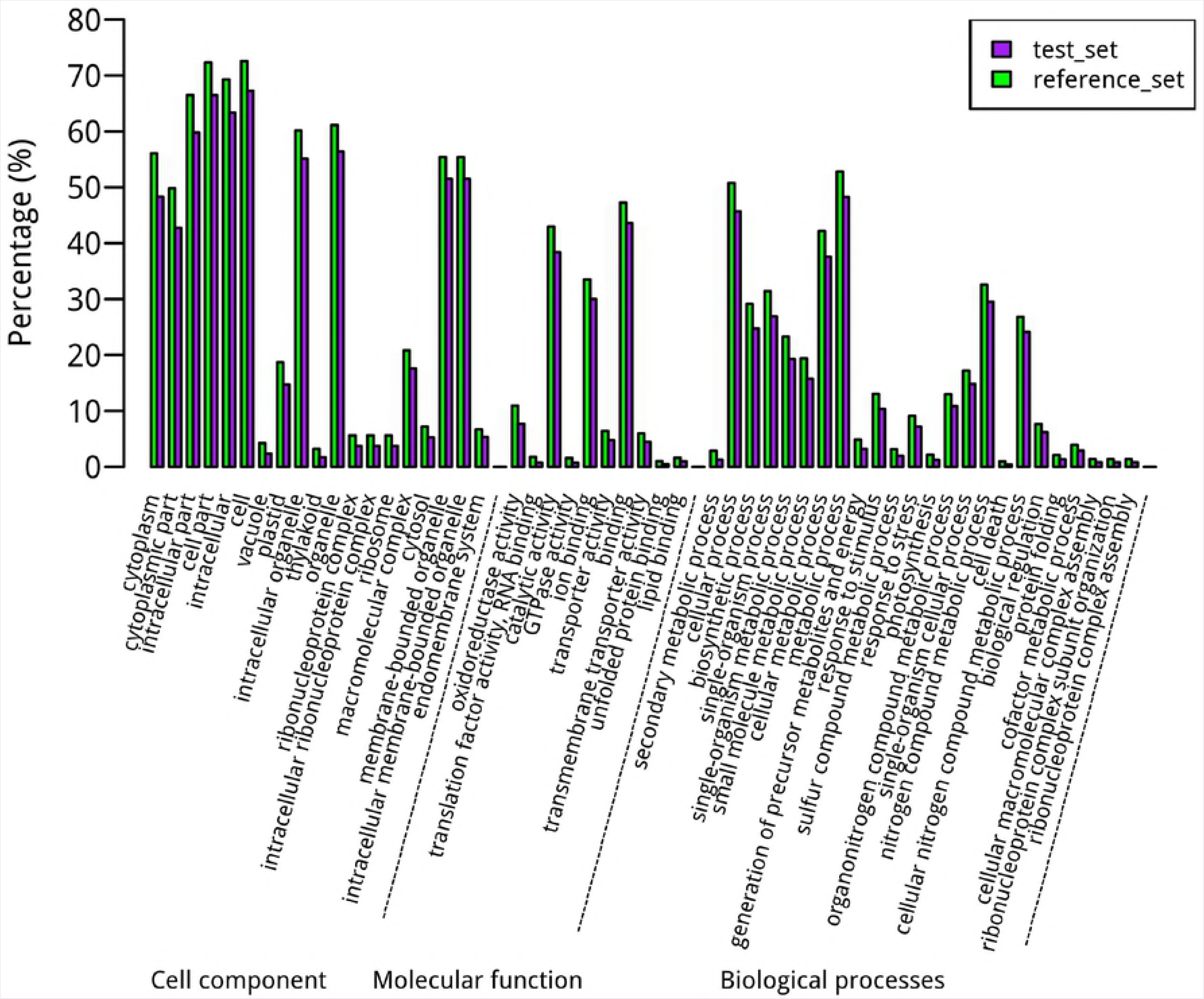
Enrichment analysis for the function of circRNAs as miRNA decoys. The GO terms containing BP (biological processes), MF (molecular functions) and CC (cell components). The GO annotation is presented on the x-axis legend and the percentage of genes on the y-axis legend (Fisher’s test, *P* < 0.05).

## Discussion

At present, the exploration of circRNA is particularly scarce in plants compared to animals. To date, circRNAs studies in plant species have been confined to several model plants. In this article, 3715 unique circRNAs were identified systematically, which can be classified into three different groups. Then the functional analysis of circRNAs was conducted and our results provide a basis for subsequent researches.

### Complementary sequences and inverted repeat pairs are not the main driving force underlying backsplicing of circRNAs

In animals the driving force underlying backsplicing of circRNAs was complementary sequences or repetitive sequences in the introns flanking back-spliced exons that form double-stranded RNA structures to bring the splice sites adjacent to one another [23, 31, 41-43]. However, it is not clear whether the intron pairing was also the predominant driving force for circRNA formation in plant. Comparing the intron length distributions of circRNAs with that of linear gene, we found that longer introns maybe involved in the formation of circRNAs in maize, which were also found in tomato (S5 Fig). So this feature of longer introns flanking circularized exons found in maize, tomato, rice and thale cress maybe a common feature of plant circRNAs [15]. In addition, a total of 0, 1, 3, 4, and 7 complementary sequences were present in the 20 nt, 50 nt, 100 nt, 200 nt and 500 nt bins of introns flanking circularized exons in maize, respectively (S9 Table). The max percentage of complementary sequences in introns was 0.19% for the total 3715 circRNAs. A total of 55.6% of the complemented sequences had short lengths less than 100 nt, and the longest length was 472 nt (S9 Table). The alignment of the intronic sequences to the repetitive sequences showed that only 14 out of 3715 circRNAs overlapped with annotated elements in the 500 bp intronic regions. However, of the 7 circRNAs with reverse complements in the 500 bp region, 2 possessed repeats that were detected only in the downstream region (i.e., with no pairing ability) (S5 Fig). In conclusion, only handful of complementary sequences and inverted repeat pairs detected in the intron flanking circRNAs, which markedly differed from the results obtained in animals and suggested that other factors or driving forces were likely to be involved in the formation of circRNAs.

### The potential function of circRNAs

To date, the functions of circRNAs are largely unknown. By using our developed methods, the function of circRNAs can be inferred. To investigate whether circRNAs acted as miRNA decoys in maize, the method used in our previous reports was used to identify circRNAs as miRNA decoys [39]. Overall, 346 unique circRNAs out of 2007 exonic circRNAs were potential decoys for 144 unique miRNAs. Interestingly, when compared circRNAs as miRNA decoys with our previous results (lincRNAs as miRNAs decoys) [39], we found that the ratio of circRNAs as miRNA decoys was higher than that of lincRNAs as miRNA decoys, which may imply their different roles in regulating gene expression.

In addition, the function of circRNAs was further predicted based on the ceRNA hypothesis and GO analysis. By comparing the potential function of the circRNAs from the parent gene and acting as miRNA decoys, the GO items (53) of the circRNAs as miRNA decoys more than the items (14) of circRNAs from the parent gene. But the functional prediction results of circRNAs from this two parts were consistent in several aspects, such as components of cytoplasmic, catalytic and binding activity, response to stress and intracellular metabolic processes. Several pathway refer to energy transformation (carbon fixation pathways, starch and surcrose metabolism, nitrogen metabolism, and photosynthesis) suggests circRNAs may contribute to crop products. Therefore, these results further prove that circRNAs could participate in a variety of biological processes, constitute different cell components and exert a variety of molecular functions.

Although the results of circRNAs’ functional prediction and expression profiling in this paper show that maize circRNAs may be involved in a variety of metabolic processes and stresses in plants, the function of plant circRNAs need to be further validated.

## Conclusions

This study found 3715 unique circRNAs in maize across different experiments by employing a computational pipeline for the genome-wide identification of circRNAs. Furthermore, the property of circRNAs, patterns of differential expression and tissue-specific expression are investigated. Finally, possible functions of circRNAs are inferred by two different methods. Our method and results provide an in-depth analysis of maize circRNAs and can be expanded to more plant species. In addition, future experimental studies are required to elucidate the mechanisms and functions of circRNAs in plants.

## Materials and Methods

### Data sets

The *Zea mays* genomes and gene annotation files were downloaded from Phytozome v9.0 (http://phytozome.jgi.doe.gov/pz/portal.html). Transcriptome data were downloaded from NCBI under accession number SRP006965 (containing embryo sac, ovule, mature pollen, and seedling), SRP011480 (only the B73 line for unpollinated ear tip, seedling shoot, immature tassel, and seedling root), SRP041183 (presenting the transcriptome of *Z. mays* genotypes under control and stress conditions), SRP052520 (representing drought stress treatment of 14-day seedlings) and SRP061631 (three tissue of leaf, root and stem under control and drought condition). The repetitive sequence data were downloaded from the Plant Repeats Database (http://plantrepeats.plantbiology.msu.edu/downloads.html). Find_circ used to identify maize circRNAs was downloaded from https://github.com/marvin-jens/find_circ.

### Computational identification and characteristics of circRNAs

The algorithm find_circ [2] was mainly used to predict circRNAs in our study. Generally, the main pipelines are as follows. Transcriptome data were transferred to the computational method (find_circ) to identify circRNAs in the genome. Concisely, all reads were mapped to the reference genome by Bowtie2 (v2.1.0)[44]. Then, 20 nucleotide sequences from both ends of the unmapped reads were aligned independently to the reference genome to find unique anchor positions. The anchors located in genomes with a reversed orientation from the initial order in the reads were regarded as circRNA splicing events. The GU/AG splice sites should be satisfied when the splicing event occurs around a breakpoint. The following additional filters were used: not less than two junction reads to support the circRNA splicing and no more than a 100 kb splicing distance in the genome. In-house perl scripts were used to analyze the characteristics of maize circRNAs.

### Conservation analysis of circRNAs

To explore circRNA conservation, circRNA candidates predicted as exonic circRNAs were selected in *Z. mays, A. thaliana* and *O. sativa* [15]. To determine the orthologous gene pairs between species, *Z. mays* protein sequences were blasted against with *A. thaliana* and and *O. sativa* protein sequences (BlastP in BLAST+, v2.2.26, E < 1e-10), and the best paired genes were selected as orthologs by using our in-house Perl scripts. Then circRNAs from orthologous parental genes in both species were reanalyzed and regarded as conserved circRNAs by using BLAST (BLAST+, v2.2.26). Another two species, *S. lycopersicum* and *G. max* were analyzed with the same method. The alignment programme mVISTA was used to globally align DNA sequences to allow identification of sequence similarities and differences (*P* < 0.05).

### Validation of maize circRNAs

Maize (Zea *mays L*.) seedlings were first grown for 2-week-old in the greenhouse (30/22°C of day/night temperature, a 16–h light/8–h dark cycle). Total RNA was isolated from leaves of the maize seedlings using TRIzol reagent (Ambion) according to the manufacturer’s instructions. Partial total RNA samples were treated for 15 min at 37°C with 3 units μg^™1^ of RNase R (Epicentre). RNase R could digest essentially linear RNAs. First-strand cDNA was synthesized from untreated and RNase R-treated total RNA using the iScript cDNA synthesis kit (Bio-Rad) respectively. Polymerase chain reaction (PCR) primers (divergent and convergent) were designed for circRNA validation (S4 Table). The reagent 2 × Taq Master Mix (Vazyme, Nanjing, CN) was used for cDNA and gDNA amplification. PCR amplification conditions were as follows: 5 min pre-denaturation at 94°C; 34 cycles of 30 s denaturing at 94°C, 40 s annealing at the suitable temperature according to the primers, 30 s extension at 72°C; 10 min extension at 72°C. Then, Sanger sequencing was performed on all PCR products.

### Expression analysis of circRNAs and their parent genes

The expression level of each circRNA was evaluated by ratio of circular-to-linear and junction read counts (RPM) [4, 33]. The expression levels of the corresponding mRNAs were determined by RPKM values (reads per kilobase of exon model per million mapped reads in the sample). The circRNAs were classified into five levels: 0-0.5, 0.5-1, 1-5, 5-10, and >10. The parent genes were classified as 0-50, 50-100, 100-500, 500-2000, and >2000. Pearson’s correlation coefficient was used to evaluate the coexpression of circRNAs and their parent genes.

The degree of tissue specificity for circRNAs was evaluated with a tissue-specific index that ranged from 0 to 1 [45]. The tissue-specific 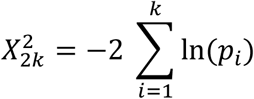, in which *n* is the number of tissues, exp_i_, is the expression value of each circRNA in tissue *i*, and exp_*max*_ is the maximum expression value of each circRNA or linear gene among 10 selected tissues. The circRNAs with an index greater than 0.9 were deemed as exhibit specific expression in one tissue. The criteria for differential expression in circRNAs and their parent genes were determined by fold changes based on the software edgeR (*P* < 0.05).

### Function predictions of circRNAs in maize

#### Functional analysis of circRNAs origined from protein-coding genes

To explore whether circRNAs were generated from parent genes with special functions, the protein sequences of the parent genes corresponding to the circRNAs were firstly blasted against the nr database, then GO enrichment and KEGG analysis were performed for the parent genes generating circRNAs by using Blast2GO with Fisher’s exact test (FDR < 0.05)[46].

#### Funcational analysis of circRNAs acting as miRNA decoys

To test whether circRNAs could act as miRNA decoys, 203 sequences of plant miRNAs downloaded from miRBase (http://mirbase.org/) were aligned to the circRNA candidate sequences using GSTAr.pl (https://github.com/MikeAxtell/GSTAr). The method used to predict miRNA decoys from circRNAs was built from previous reports [38, 39]. The general criteria were as follows: 1) one to six mismatched or inserted bases were allowed between the ninth to twentieth nucleotides from the miRNA 5’ end; 2) the position from the second to the eighth nucleotide of the 5’ end of the miRNA sequence was requested to have perfect matching; 3) no more than 4 mismatches or indels were allowed in other regions.

We also constructed and visualized the circRNA-miRNA-mRNA network by integrating miRNAs, circRNAs acting as miRNA decoys and mRNAs acting as miRNA targets [39]. The functions of the circRNAs acting as miRNA decoys were speculated based on the circRNA-miRNA-mRNA networks according to the ceRNA hypothesis and GO analysis. The IDs of all listed mRNAs connected with circRNAs acting as miRNA decoys were firstly submitted to Blast2GO, then conduct the GO enrichment analysis using Fisher’s exact test (FDR < 0.05).

#### The driving force underlying backsplicing of circRNAs

According to the annotation file, the different gradient intronic sequences flanking the backsplice sites (20 nt, 50 nt, 100 nt, 200 nt, and 500 nt) were extracted. To test the extent of reverse complementation of intronic sequences flanking circRNAs, those sequences used as input and database were blasted to itself (BlastN in BLAST+, v2.2.26, E < 1e-5, word size 5). The BLAST results were examined using our custom Perl scripts. In addition, repetitive sequences were aligned to 500 nt downstream and upstream of intronic sequences flanking the backsplice sites to determine whether the repetitive elements played a role in the circularization mechanism.

## Supporting information

### S1 Fig. Workflow used to predict circRNAs

Schematic diagram for the prediction of circRNAs in accordance with the method in a previous report.

### S2 Fig. Alternative circularization and conservation

(A) Example of single and multiple circularized exon and type of alternative circularization. Backspliced reads are marked. Orange rectangles: exons in transcripts; yellow rectangles: exons in another transcript derived from the same gene in the third example. Black line at the exon boundaries: backsplice site overlapped with known exon boundaries; red line: novel splice site. Absolute green arrows: backsplice using both annotated splice sites; green arrows with a blue rim: backsplice using two annotated splice sites from different transcripts; blue arrows: backsplice using no known exon boundaries. (B) Another example of sequence conservation analysis of circRNAs between *A. thaliana* and *Z. mays*. The label is the same as Fig 2C.

### S3 Fig. Expression profiling of circRNAs and their parent genes

Related to Fig 3. (A) Expression level of circRNAs measured by the ratio of circular to linear in different scopes. (B) Expression level of parent genes overlapped with circRNAs in different ranges. (C) Heatmap of circRNAs with tissue-specific expression. (D) Differential expression of circRNAs and their parent genes under heat conditions in the seedlings.

### S4 Fig. Functional analysis of circRNAs

(A) Enrichment analysis for genes overlapped with circRNAs that were significant (Fisher’s test, *P* < 0.05). The x- and y-axes are the same as Figure 5. (B) Genome-wide miRNA-regulated networks. Pink nodes: circRNAs; blue nodes: miRNAs; green nodes: mRNAs. Grey edges: correlations.

### S5 Fig. Feature of intron flanking circularized exons

(A) Mean length distribution of upstream and downstream intron flanking circularized exons in *tomato*. (B) Example for intron pairing of reverse complementation in flanking sequences of the backsplice sites. Repetitive elements exist in the downstream of these complementary sequences (red colour). A part of the pair of the second circRNA is shown.

S1 Table. Prediction of circRNAs from various transcriptome data sets in *Z. mays*.

S2 Table. Alternative circularization phenomenon and whether the backsplice site is annotated.

### S3 Table. Result of Conservation analysis

(A) Result of Conservation analysis between maize and rice. (B) Result of Conservation analysis between maize and *Arabidopsis Thaliana*. (C) Result of Conservation analysis between maize and Soybean. (D) Result of Conservation analysis between maize and tomato.

### S4 Table. Primers used to validate maize circRNAs

(A) Divergent primers for validation of candidate circRNAs. (B) Convergent primers for validation of candidate circRNAs.

S5 Table. Correlation of circRNAs and their corresponding parent genes.

### S6 Table. Functional prediction of circRNAs originated from protein-coding genes indirectly

(A) CircRNAs and their parent genes function. (B) Pathway statistics

### S7 Table. List of circRNAs acting as miRNA decoys

miRNA were listed in the first column. CircRNA in the second column acting as the miRNA decoys. The following columns mean the starting and terminating sites between circRNA and miRNA, MFE of a perfectly matched site, MFE of the alignments, MFEsite/MFEperfect, respectively. MFE: minimum free energy.

S8 Table. Gene ontology (GO) analysis of circRNAs acting as miRNA decoys based on ceRNA hypothesis.

### S9 Table. Reverse complement in different intervals of intron flanking circularized exons

The length was extracted in 50 nt, 100 nt, 200 nt and 500 nt intronic regions.

## Abbreviations

circRNAs: Circular RNAs
RPM: Reads per million mapped reads
ss: Splicing site
miRNAs: microRNAs
RBP: RNA binding proteins
ecircRNA: Exonic circRNAs
CDS: Coding sequence
UTR: Untranslated regions
AS: Alternative splicing
RPKM: Reads per kilobase of exon model per million mapped reads
IRES: Internal ribosome entry site
ORF: Open reading frame
ceRNA: Competing endogenous RNAs
GO: Gene ontology
lincRNAs: Long intergenic noncoding RNAs
nt: Nucleotide
dr: Drought
ctrl: Control

## Acknowledgements

This work was supported by grants from the National Natural Science Foundation of China (Grant No.31070256, No.31370329 and No.11631012), the Program for New Century Excellent Talents in University (NCET-12-0896), the Natural Science Basic Research Plan of Shaanxi Province, China (No. 2014JM3074), and the Fundamental Research Funds for the Central Universities (No. GK201403004). The funding agencies had no role in the study, its design, the data collection and analysis, the decision to publish, or the preparation of the manuscript.

## Author Contributions

**Conceptualization:** Baihua Tang, Guanglin Li.

**Data curation:** Baihua Tang, Zhiqiang Hao.

**Formal analysis:** Baihua Tang, Zhiqiang Hao, Yanfeng Zhu, Hua Zhang.

**Writing - original draft:** Baihua Tang, Zhiqiang Hao, Yanfeng Zhu, Guanglin Li.

**Writing - review & editing:** Baihua Tang, Zhiqiang Hao, Yanfeng Zhu.

